# Crystallization of magnesium calcite otoconia in the inner ear of the developing quail

**DOI:** 10.64898/2026.08.10.743921

**Authors:** Einat Kedar, Jia Hui Lim, Ernesto Scoppola, Peter Fratzl, Shahrouz Amini, Emeline Raguin

**Affiliations:** Department of Biomaterials, Max Planck Institute of Colloids and Interfaces, Potsdam, Germany; Department of Sustainable and Bio-inspired Materials, Max Planck Institute of Colloids and Interfaces, Potsdam, Germany

**Keywords:** Avian otolith, Otoconia nanostructure, Magnesium-Calcite, Lagena development, Polycrystalline, Crystallography

## Abstract

Otolith organs are specialized structures of the vertebrate inner ear that provide the inertial mass required for maintaining equilibrium. Mammals possess two otolithic organs, the utricle and the saccule, whereas birds and other non-mammalian vertebrates also retain a third one, the lagena, whose development remains poorly understood and whose function is still debated. In birds, the lagena contains thousands of calcium carbonate biomineral particles, termed otoconia. Here, we reconstruct the developmental crystallization of lagena otoconia in the Japanese quail (*Coturnix japonica*) throughout embryogenesis. We combine multiscale imaging with structural, compositional, and crystallographic analyses across length scales. We show that lagena mineralization precedes cranial bone formation and proceeds predominantly through the growth of existing otoconia rather than continued nucleation. Otoconia develop through progressive particle growth, alignment, and fusion within a pre-existing organic compartment while maintaining a persistent central core, ultimately forming magnesium calcite biominerals. This maturation is accompanied by progressive nanoscale densification, transforming early mineral deposits into mature hierarchical crystals. This work establishes a developmental model of avian otoconia formation and provides new insights into how hierarchical calcium carbonate crystals are assembled during vertebrate development.

**Statement of significance:** Otoliths are the only calcite based biomineral in our body that has a physiological function, yet their developmental assembly remain incompletely understood. While the utricle and saccule have been extensively investigated across vertebrates, the lagena, a third otolithic organ lost during mammalian evolution, has received comparatively little attention. Here, we combine multiscale imaging and materials characterization to reconstruct the developmental crystallization of lagena otoconia in the Japanese quail. We establish how hierarchical magnesium calcite biominerals emerge through coordinated mineral growth, structural maturation, and crystallization, providing a developmental framework for avian otoconia formation and new insights into the assembly of vertebrate calcium carbonate crystals.

## 1. Introduction

Otoliths organs are specialized vestibular structures of the vertebrate inner ear that responsible for maintaining equilibrium by detecting linear acceleration and gravitational forces. Human and most mammals possess two otolith organs, the utricle and the saccule, which are sensitive to horizontal and vertical accelerations, respectively [1]. In contrast, fish, amphibians, reptiles, birds, and monotremes, possess a third otolith organ, the lagena [2]. Like the utricle and saccule, the lagena consist of mineralized particles embedded within a gelatinous otolithic membrane that overlies the sensory epithelium of the macula. In these vestibular organs, the minerals provide the inertial mass required to convert gravitational and acceleration forces into mechanical stimulation of underlying sensory hair cells [1–4]. In fish, each otolith organ contains a single mineralized otolith, whereas in birds and other tetrapods, it consists of thousands of individual biominerals known as otoconia [3, 5, 6].

The otoconium is a barrel-shaped biomineral composed of organic and inorganic components. Otoconia are composed primarily of calcium carbonate. Depending on the species and vestibular organ, this mineral phase may occur as calcite, aragonite, or, more rarely vaterite [3, 6]. Mature otoconia exhibit a complex hierarchical organization in which organic fibrillar networks are integrated within the mineral phase [7]. Mineral density is not uniformly distributed throughout the otoconium, with denser mineralization occurring at the rhombohedral facets and peripheral regions, whereas the central body is comparatively porous [6–12]. Studies in mammalian have revealed the presence of an organic core that becomes exposed following decalcification [5] and age-related degeneration [7, 13]. This organic compartment is thought to play a central role in otoconium formation by providing a scaffold that regulates crystal nucleation and growth [4–6, 8, 14].

Although the composition and structure of mature otoconia have been extensively characterized, the developmental processes leading to their formation remain poorly understood. Studies of the utricle and saccule have shown that otoconia formation is a highly orchestrated developmental process [4]. During embryogenesis, otoconia first appear as small, mineralized structures and progressively acquire their characteristic morphology as they increase in size [8–10, 15–18]. Regional variations in otoconia size and abundance have been observed across the sensory epithelium [1, 10, 18, 19], suggesting that otoconia formation is spatially regulated and may contribute to the functional specialization of vestibular organs. However, these observations are derived almost exclusively from studies of the utricle and saccule, and it remains unknown whether similar developmental and organizational principles apply to the lagena.

Furthermore, unlike the utricle and saccule, whose organization is broadly conserved across vertebrates, the lagena exhibits substantial variation in morphology across the vertebrate groups that possess one. Lagena differences have been reported in its size, shape, anatomical location, orientation, and structure [2, 20]. Interestingly, despite the remarkable diversity of avian species in terms of ecology, behavior, and locomotion, the overall organization of the lagena remains relatively similar across birds. This combination of morphological variability across vertebrates and relative conservation within birds suggests that specific structural features of the avian lagena may reflect functional adaptations unique to this organ [2].

Early studies have excluded a major auditory function in birds, [21–24]. Alternative hypotheses, including a possible involvement in magnetoreception, have been proposed but remain controversial [25–29]. Nevertheless, projections from the lagena terminate in vestibular nuclei, supporting a role in balance and spatial orientation [24]. However, previous investigations of the avian lagena have relied largely on two-dimensional observations, providing limited information on the three-dimensional (3D) growth, ultrastructural organization, crystallization pathway, and mineral composition of its otoconia.

Here, we combine multiscale 3D imaging with ultrastructural and spectroscopic analyses to characterize the growth, ultrastructure, crystallization pathway, and mineral composition of lagena otoconia during embryonic development in the Japanese quail (*Coturnix japonica)*. This approach enables us to establish a developmental model of otoconia formation from the earliest mineral deposits to the mature magnesium calcite crystal.

## 2. Materials and Methods

### 2.1 Sample preparation

In accordance with the German Animal Welfare Act and the Laboratory Animal Welfare Ordinance, no approval by an ethics committee for animal experimentation was required. Fertilized Japanese Quail eggs (*Coturnix japonica*) were purchased from commercial breeders (Wachtel-Shop Michael Volk e.K., Obersulm, Germany). Upon delivery, eggshells were gently wiped with 70% ethanol to minimize microbial contamination, placed in an automatic digital egg incubator (Ovation 56 eco egg incubator, Brinsea, North Somerset, UK) at 38.0 ± 0.5 °C and 50 ± 5% relative humidity, and automatically turned for 15 minutes every hour.

Embryos were sacrificed by cervical dislocation between embryonic developmental day (EDD) 7 and 13. For each developmental stage, at least three skulls were immersed overnight in 2% glutaraldehyde in phosphate buffered saline (PBS), whereas the remaining specimens were fixed in 4% paraformaldehyde (PFA) in PBS. All skulls were then kept in 70% ethanol for at least two days prior to further processing. For each embryonic day, several lagenae were extracted according to the procedure described in supplementary S1.1.

For microcomputed tomography (micro-CT), one skull from each developmental stage was stained with 0.1% ruthenium red following the protocol of Gabner et al. [30] and embedded in polymethyl methacrylate (PMMA; Technovit 9100, Kulzer, Germany) according to Moreno Jiménez et al. [31]. For histological analyses, the complete inner ear was dissected from the temporal bone following Otto et al. [32] and embedded in PMMA.

For transmission electron microscopy (TEM), synchrotron X-ray fluorescence microscopy (XRF), synchrotron X-ray diffraction (XRD), and energy dispersive X-ray spectroscopy (SEM-EDS), lagenae were extracted from glutaraldehyde fixed specimens and embedded in epoxy resin (EMbed 812 Embedding Kit, Electron Microscopy Sciences) following the procedure of Luft (1961)[33]. For SEM-EDS, two embedded blocks were cross-sectioned according to the procedure describe in supplementary (S1.2).

### 2.2 Structural characterization

PMMA embedded inner ears were sectioned at 6 µm thickness using a Leica microtome (Leica Biosystems Nussloch GmbH, Nussloch, Germany). Sections were stained with a Hematoxylin and Eosin Rapid Kit (Clin Tech Ltd, Guilford, UK) following the manufacturer’s instructions. Histological sections were examined using a Keyence VHX 5000 digital microscope (Keyence Corp., Osaka, Japan). Polarized transmitted light microscopy (Zeiss AXIOLAB 5) was used to evaluate the birefringence and crystallographic orientation of individual otoconia.

Whole stained skulls, PMMA embedded inner ears, and extracted otoconia were scanned using an EasyTom 150/160 micro-CT system (RX Solutions, Chavanod, France). For scans of extracted otoconia, the nanotube X-ray source was operated at 70 kV and 140 µA with a voxel size of 0.50 µm. Three-dimensional reconstructions were generated from projection images using X Act software (RX Solutions, Chavanod, France).

For scanning electron microscopy (SEM), extracted otoconia were mounted on carbon adhesive tape and coated with a 10 nm carbon layer. Samples were imaged using a Quattro S (ThermoFisher Scientific) environmental scanning electron microscope (FEI, Hillsboro, OR, USA) equipped with a backscattered electron detector operating in low vacuum mode (0.5-1 Torr), with accelerating voltages ranging from 5 to 12.5 kV.

For transmission electron microscopy, epoxy embedded lagena were sectioned using an RMC Boeckeler PowerTome ultramicrotome (Arizona, USA). Ultrathin sections of 70, 100, and 120 nm thickness were mounted on 200-mesh copper grids. TEM characterization was performed using a JEOL JEM-F200 microscope (JEOL Ltd., Japan) operating at 200 kV and equipped with a cold field-emission gun (cFEG). Bright-field images were recorded on a Gatan Metro 2k × 2k direct electron detector (Gatan Inc., USA) using Gatan Digital Micrograph software. Selected-area electron diffraction (SAED) was used to determine the crystallographic phase and orientation of the otoconia. Dark-field images were formed by selecting the calcite (104) reflection to visualize crystallographic orientation and reveal misoriented domains. The images and diffraction patterns were analyzed using the Fiji program.

For focused ion beam scanning electron microscopy (FIB-SEM), extracted otoconia (PFA fixed) from each developmental stages investigated here, were transferred onto aluminum SEM stubs and coated with 10nm carbon and 10nm platinum. Samples were imaged using a Crossbeam 540 FIB-SEM (Zeiss, Oberkochen, Germany). Serial sectioning was performed at 30 kV and 100 pA, with the exposed surface imaged after each milling step at 2 kV and 500 pA. Images were acquired simultaneously using the dual channel mode, combining mixed InLens and secondary electron (InLens/SE) and backscattered electron (BSE) signals. While the InLens/SE detector provided detailed surface topography, the BSE detector generated atomic number contrast, enhancing the visualization of denser mineralized regions within the otoconia.

### 2.3 Compositional characterization

Elemental composition of individual otoconia was investigated by energy dispersive X-ray spectroscopy (EDX) using the Quattro S (Thermo Fisher Scientific, FEI Deutschland GmbH, Germany) environmental scanning electron microscope. Analyses were performed in single point acquisition mode under high vacuum conditions with an accelerating voltage of 15 kV, a spot size of 5.0, and a lifetime limit of 80. To achieve conductivity, sample were coated with 10nm carbon layer. Additional area analysis was performed on a FIB cross-section block. The block was mounted flat onto an SEM stub using conductive carbon tape. The sample was coated with a 20 nm carbon layer prior to analysis. EDS measurements were carried out in the ZEISS Crossbeam 550 using an Ultim Max 170 EDS detector (Oxford Instruments, United Kingdom). The EDS map was obtained with electron beams of an accelerating voltage of 10 kV and emission current of 200 uA.

Raman spectroscopy was performed to characterize the mineral composition of otoconia. Single point spectra were acquired from an EDD13 otoconium and compared with geological calcite and quail eggshell measured under identical conditions. Measurements were performed using a WITec alpha300R confocal Raman microscope equipped with a 532 nm excitation laser and a Zeiss EC Epiplan Neofluar DIC 20× objective (NA = 0.5). Spectra were acquired using an integration time of 1 s, 45 accumulations, and a laser power of 15 mW.

Synchrotron X-ray fluorescence microscopy (XRF) and X-ray diffraction (XRD) measurements were performed at the mySpot beamline of the BESSY II synchrotron radiation source, Helmholtz-Zentrum Berlin, Germany [34]. The beamline was operated at an incident X-ray energy of 18 keV using the Mo/B₄C multilayer monochromator. The X-ray beam was defined by a sequence of pinholes, resulting in a beam size of approximately 30 × 30 μm² at the sample position. Samples were mounted on dedicated sample holders and raster-scanned in two dimensions with a step size of 30 μm and an exposure time of 7 s per scan point.

At each scan position, two-dimensional XRD patterns were collected using a large-area Dectris EIGER 9M detector. XRF spectra were recorded in transmission geometry using a RAYSPEC Sirius SD-E65133-BE-INC detector equipped with an 8 μm beryllium window. The XRF detector was positioned on the transmission side of the sample at approximately 50° with respect to the direct beam. The incident beam intensity was monitored for each measurement using an upstream ionization chamber. Sample transmission was estimated from the X-ray fluorescence signal emitted by a lead beamstop positioned in front of the XRD detector and collected by the XRF detector.

The detector geometry, including sample-to-detector distance, beam center, and detector orientation, was calibrated using a silicon standard, NIST SRM640. The calibrated sample-to-detector distance was approximately 350 mm. The scattering vector was defined as:

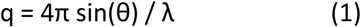

where 2θ is the scattering angle and λ is the X-ray wavelength. The scattering intensity was normalized against a glassy carbon standard [35]. Background measurements were acquired without sample for 20 s in order to improve the signal-to-noise ratio of the background-subtracted data. XRF and XRD data were corrected for background and normalized to exposure time, incident beam intensity, and sample transmission. Data reduction was performed using in-house Python routines based on pyFAI [36].

The corrected two-dimensional XRD patterns were azimuthally integrated to obtain one-dimensional scattering profiles, I(q), over a momentum-transfer range of approximately 0.1–42 nm⁻¹. XRF elemental maps were obtained by integrating the background-corrected fluorescence intensity over selected emission-line energy windows. The resulting normalized XRF intensity maps were used to assess relative elemental distributions across the samples and to correlate local composition with structural information from XRD/SAXS (small-angle X-ray scattering).

Calcite-related diffraction peaks were analyzed in terms of peak position. Lattice parameters were extracted by combining two or more calcite reflections and assuming a hexagonal lattice, yielding spatially resolved values of the a- and c-lattice parameters.

The small-angle scattering region between 0.1 and 4 nm⁻¹ was analyzed using a T-like parameter approach inspired by the T-parameter analysis commonly applied to mineralized tissues [37, 38]. In classical mineralized platelet systems, the T-parameter represents a characteristic mineral thickness. Here, however, because otoconia do not correspond to this geometry, the calculated parameter was not interpreted as an absolute particle thickness. Instead, it was used as a relative characteristic length describing developmental variations in the nanoscale particle/interface organization and is referred to throughout the manuscript as the apparent pore size.

After background subtraction and normalization, the Porod constant, P, was obtained by assuming a Porod slope of −4 in the high-q part of the selected SAXS region, corresponding to scattering from sharp interfaces. In practice, P was calculated from the Porod limit:

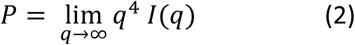

within the q-range where the Porod regime was observed. The integrated SAXS intensity, J, was calculated by integrating the scattering curve in Kratky representation over the selected small-angle range:

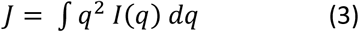

The apparent T-like parameter was then calculated as:

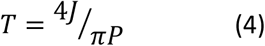

Accordingly, the resulting values should be considered apparent characteristic length scales associated with particle and pore interfaces rather than direct measurements of physical pore dimensions.

### 2.4 Image processing, segmentation and data analysis

All micro-CT images were processed by Dragonfly software Version 2024.1 (Object Research Systems (ORS) Inc, Montreal, Canada). The cartilaginous labyrinth, bone and otoconia were manually segmented using a combination of brush and intensity threshold selection. Otoconia were individualized using the watershed module. Volume measurements and volume distribution maps were generated using the compute measurements and measurements inspector modules. Otoconia count and volume measurements were analyzed via RStudio script.

FIB-SEM stacks were aligned using the Sum of Squared Differences (SSD) registration algorithm in Dragonfly software (Version 2024.1; Object Research Systems (ORS) Inc., Montreal, Canada). Mixed InLens/secondary electron datasets were enhanced using a convolution filter followed by Contrast Limited Adaptive Histogram Equalization (CLAHE) to improve signal to noise ratio and local contrast. Backscattered electron datasets were processed using a convolution filter only.

## 3. Results

### 3.1 Morphology and growth of the skull, the inner ear and the lagena otolith in quails

The anatomical position of the quail inner ear at EDD11 is shown in Fig. 1A. The lagena is located at the distal end of the basilar papilla and exhibit a characteristic U-shaped morphology (Fig. 1A-B).

**Fig. 1.**
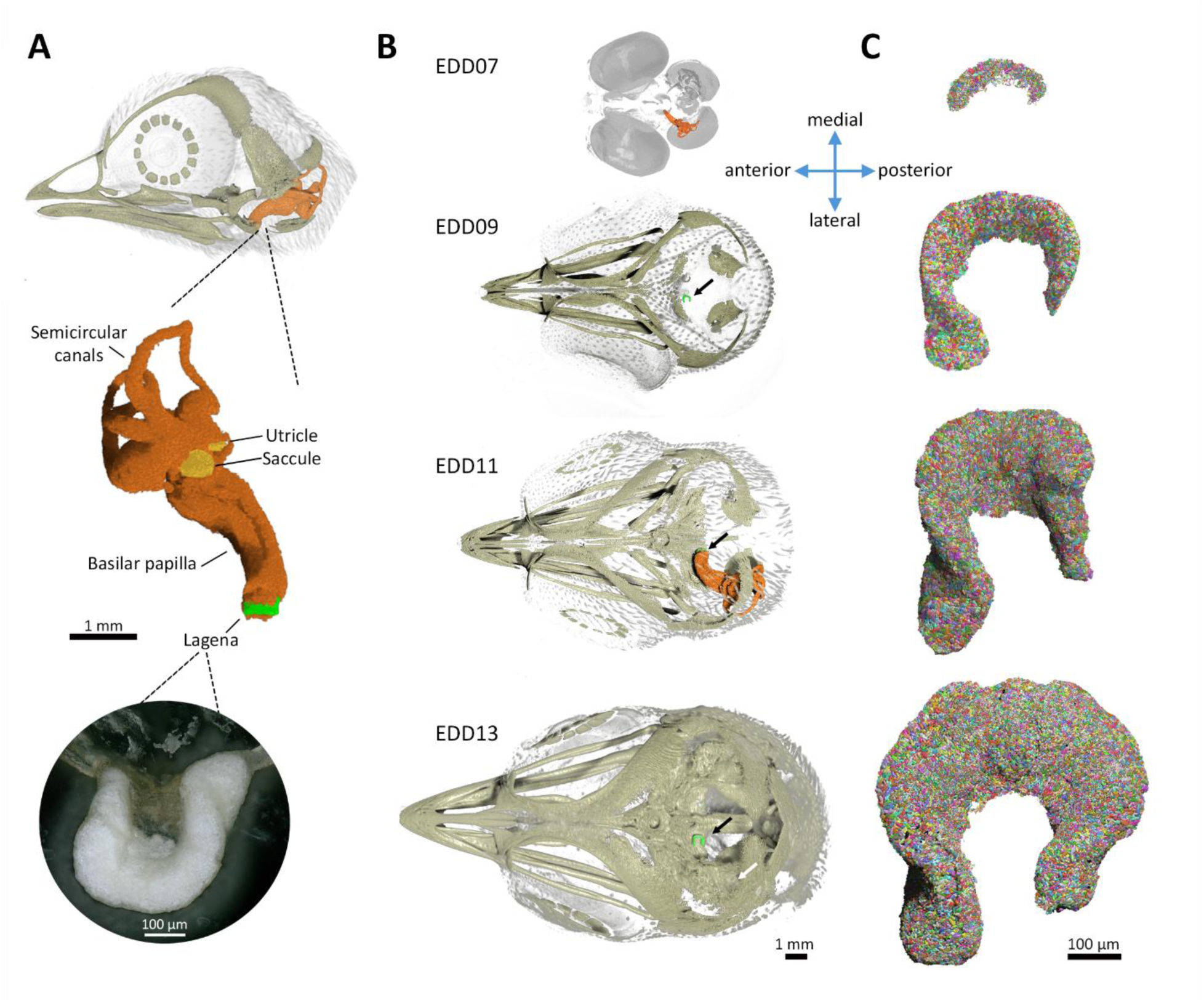
The skull, inner ear and lagena, at different embryonic development days (EDD). (A) Left lateral view of the skull at EDD11: the anatomic position of the left cartilaginous labyrinth (orange). Medial view of the inner ear (enlarged): the utricle and saccule (yellow) located on the vestibule are oriented parallel to the horizontal and vertical planes, respectively, and the lagena (green) is located at the end of the basilar papilla. Light microscopy image of the lagena at EDD11 (700x), see supplementary for enlarge image (Fig. S2.1). (B) Dorsal (top) view of the skull: at EDD07 the soft tissue (in gray) is visible, and the cartilaginous labyrinth is fully constructed (orange). The black arrows point on the lagena location in the skull mid-sagittal line at EDD09, at EDD11 (also in relation to the inner ear, in orange) and at EDD13. The white arrow at EDD13 shows that the bony labyrinth is almost completed, covering most of the inner ear. (C) Dorsal (top) view of left lagena, extracted and scanned in high resolution (0.5 µm voxel size), demonstrates its development. EDD titles and orientation map corresponded to B and C.

The growth of the skull and lagena throughout embryonic development is presented in Fig. 1B-C. At EDD07, the earliest developmental stage examined here, the cartilaginous labyrinth is fully formed, while no bone mineralization is seen (Fig. 1B). In contrast, mineral deposits are already present within the lagena (Fig. 1C), suggesting that lagena mineralization precedes skeletal mineralization. At EDD09, bone mineralization is in progress and the lagena, visible near the mid sagittal plane of the skull (Fig. 1B-C: EDD09 black arrow), had expanded substantially and acquired its characteristic shape, in which the anterior end is longer than the posterior one. At EDD11, growth occurs in the central region of the lagena, resulting in a marked widening of the structure. At EDD13, the lagena reaches its mature U-shape morphology, with both extremities folded inward (Fig.1C). At this stage, the bony labyrinth covers most of the cartilaginous part (Fig.1B: EDD13).

### 3.2 Otoconia interaction with the macula and crystal orientation

The interaction between otoconia and the surrounding tissues is observed in the histological section at EDD09 (Fig. 2A). Otoconia (purple) display a broad size distribution throughout the otolithic membrane. Polarized light microscopy analysis confirms the birefringence of the otoconia (Fig. 2B). Individual otoconia show changes in brightness and color upon sample rotation, while neighboring otoconia display distinct optical responses. These observations demonstrate that neighboring otoconia exhibit distinct crystallographic orientations.

**Fig. 2.**
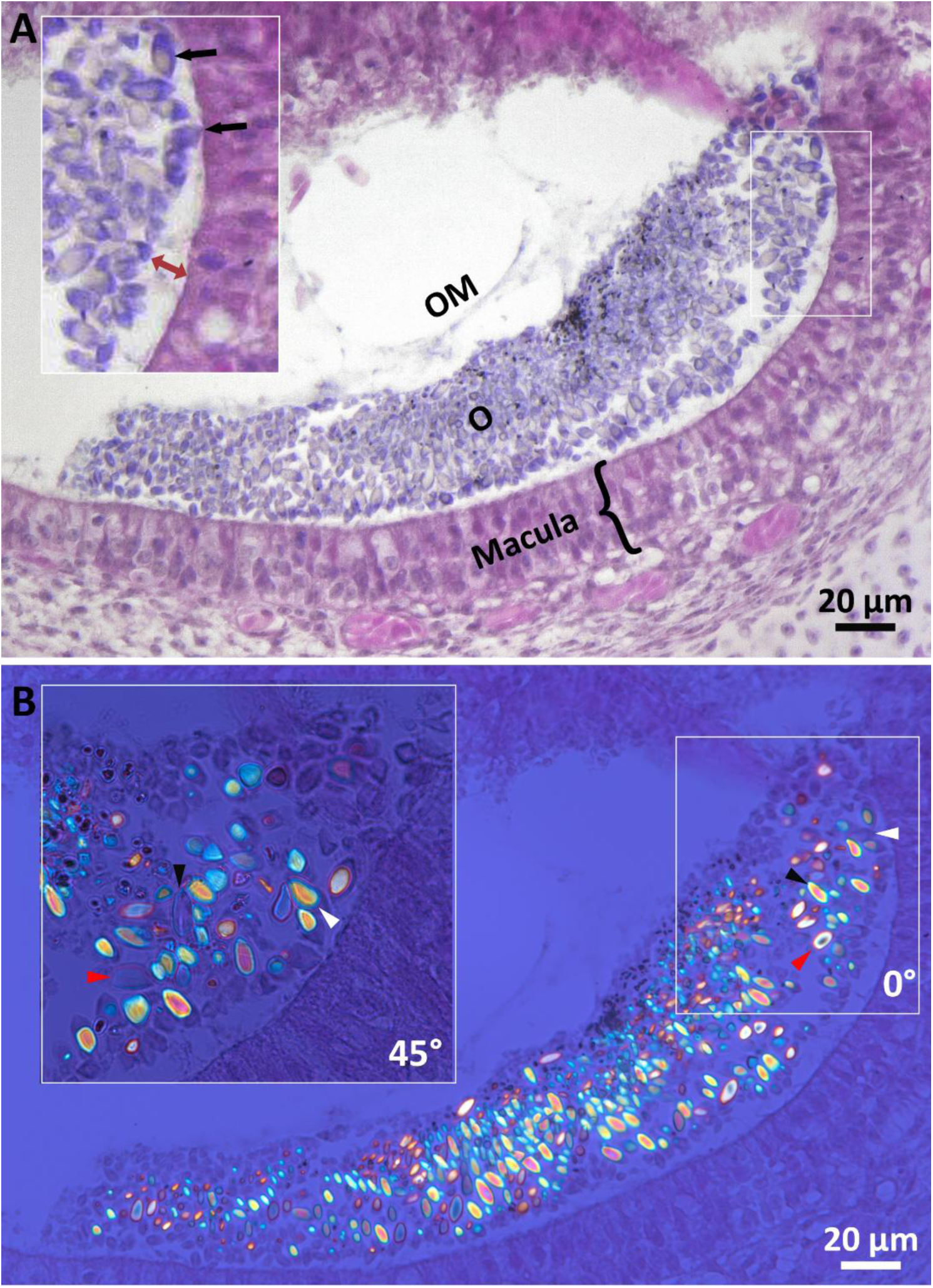
Otoconia interaction with the macula. (A) Histology section of the lagena at EDD09, 6 µm slice thickness, stained with hematoxylin and eosin. Otoconia (O) in purple, embedded within the otolithic membrane (OM), above the macule (containing the sensory cells). Inset: the gelatine membrane gap in which the cilia located (red arrow), and some otoconia (black arrows) that are in close contact with the macula. (B) Polarized transmitted light microscopy: otoconia display polarized effect as crystals reflect different colors in different directions. The change of color, up to full extinction appear when sample is turned in 45° (inset: 100x, white, red and black arrow heads each shows brightness changes of a single otoconium at each angle).

### 3.3 Growth rate of the lagena and otoconia

Micro-CT data analysis showed that both total otoconia volume and otoconia number increase markedly during development (Table 1). The most pronounced growth occurs between EDD07 to EDD09, during which total otoconia volume increases by approximately 27-fold, whereas the total number of otoconia increases approximately 15-fold. At EDD13, total otoconia volume is 130 times greater than at EDD07, while otoconia number increases about 45-fold. Accordingly, otoconia density (number of otoconia per 1 µm^3^) progressively decreases through the developmental window investigated (Table 1), indicating that enlargement of existing otoconia contributes more to the increase in total mineralized volume than the formation of new otoconia.

**Table 1.**
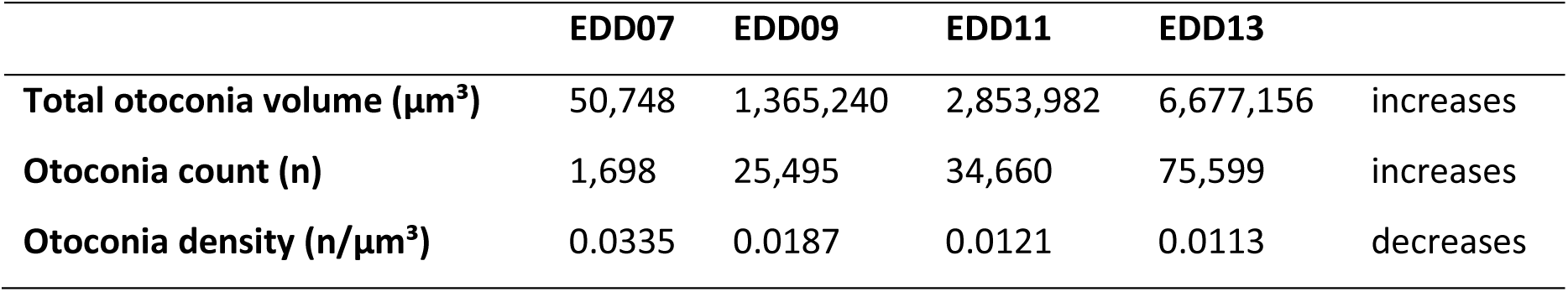
Otoconia count and total volume, by embryonic development day (EDD).

|  | **EDD07** | **EDD09** | **EDD11** | **EDD13** |  |
| --- | --- | --- | --- | --- | --- |
| **Total otoconia volume ($\mu\text{m}^3$)** | 50,748 | 1,365,240 | 2,853,982 | 6,677,156 | increases |
| **Otoconia count (n)** | 1,698 | 25,495 | 34,660 | 75,599 | increases |
| **Otoconia density (n/$\mu\text{m}^3$)** | 0.0335 | 0.0187 | 0.0121 | 0.0113 | decreases |

Analysis of individual otoconium volume supports this observation (Fig. 3A-B): at EDD07, otoconia smaller than 20 µm^3^ represent a bit less than 70% of the population but account for only 16% of the total otoconia volume. However, at this stage, grayscale intensity thresholding was limited because the early mineral particles were below the spatial resolution of the micro-CT (0.5 µm voxel size). Consequently, only larger mineral aggregates, which constituted most of the lagena mass, could be segmented. From EDD09 onwards, this population progressively decreases, representing approximately 31%, 24% and 15% of the otoconia population at EDD09, EDD11, and EDD13, respectively, while contributing less than 7% of the total mineralized volume (Fig. 3B). Conversely, larger otoconia account for an increasing proportion of the total volume throughout development. Despite these changes, the overall size distribution remains broad at all developmental stages, with no dominant size class emerging. The continued presence of small otoconia throughout embryogenesis is consistent with the ongoing formation of new mineralized particles while previously formed otoconia continue to grow.

**Fig. 3.**
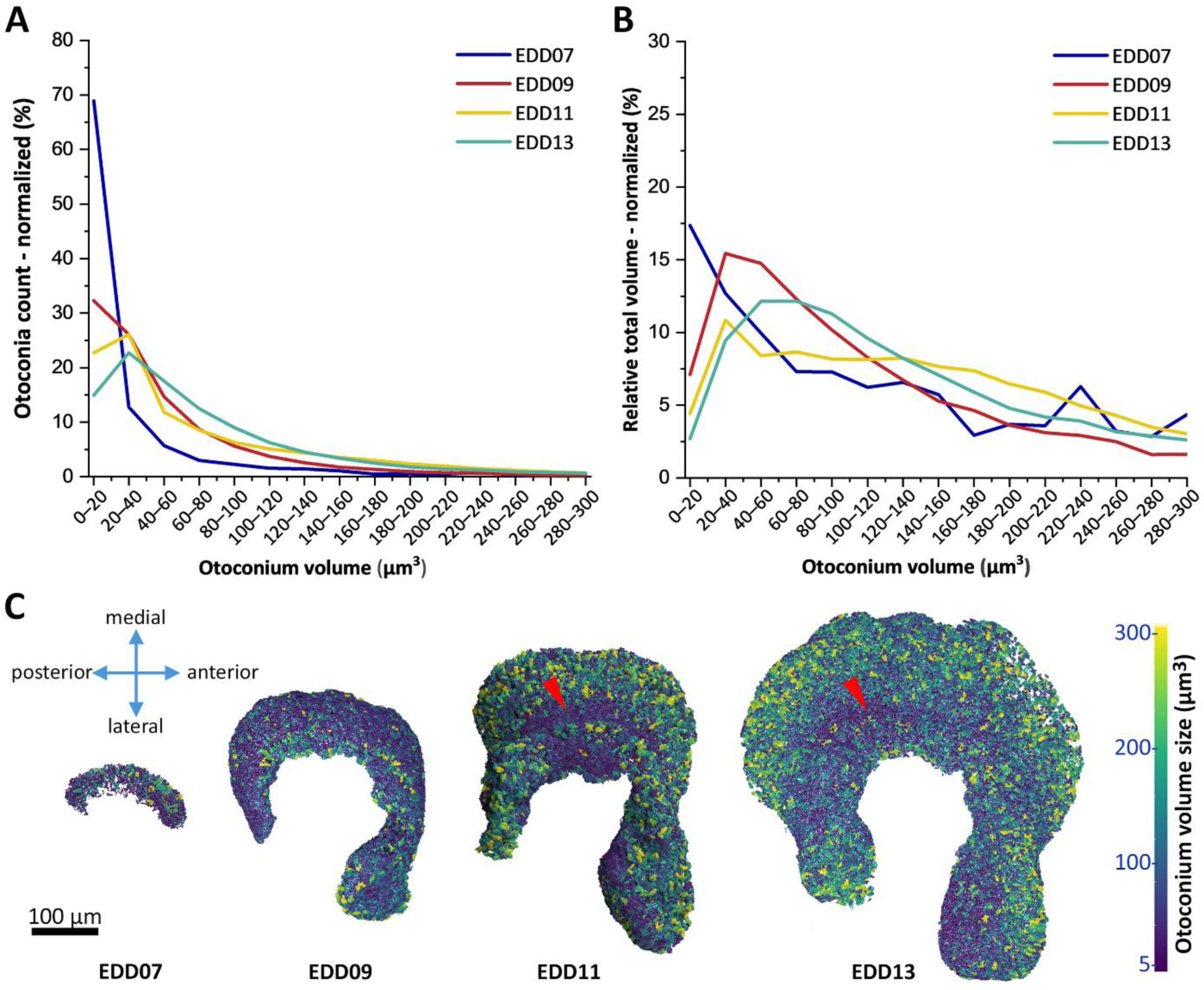
Otoconia size distribution and growth rate. (A) Size distribution of individual otoconium and (B) their contribution to the total otoconia volume. (C) Ventral (inferior) view of the lagena: otoconia distribution according to their size (between 5-300 µm^3^). Otoconia >300 µm^3^ were excluded (included portions of fused otoconia that could not be separated by segmentation).

3D micro-CT segmentation mapping further reveals a distinct spatial organization of otoconia within the lagena (Fig. 3C). At EDD07, relatively large, mineralized structures are mainly located along the periphery of the lagena. As development proceeds, the largest otoconia remain preferentially distributed near the outer surface of the membrane (green-yellow), whereas smaller otoconia become concentrated within the central region (blue-purple). This spatial organization is particularly evident at EDD11 and EDD13, where a continuous band of small otoconia is observed above the striola, the central region of the macula where hair cell orientation reverses (Fig. 3C, red arrowheads).

### 3.4 Otoconium morphology and mineralization

SEM observation reveals progressive morphological maturation of individual otoconia throughout embryogenesis (Fig. 4A-D). At EDD07, otoconia are plate-shaped mineralized units, approximately 2 µm in length (Fig. 4A). At EDD09, the otoconium grow in length (10 µm) and width (about 4 µm), and its morphology changes to barrel shaped belly with rhombohedral facets ends (Fig. 4B). From this stage onwards the shape of the otoconia remain similar, although some facets appear more pronounced at the extremities at EDD13 (Fig. 4C-D). We also noticed few very large otoconia (>30µm) at EDD13 (Fig. S2.2).

**Fig. 4.**
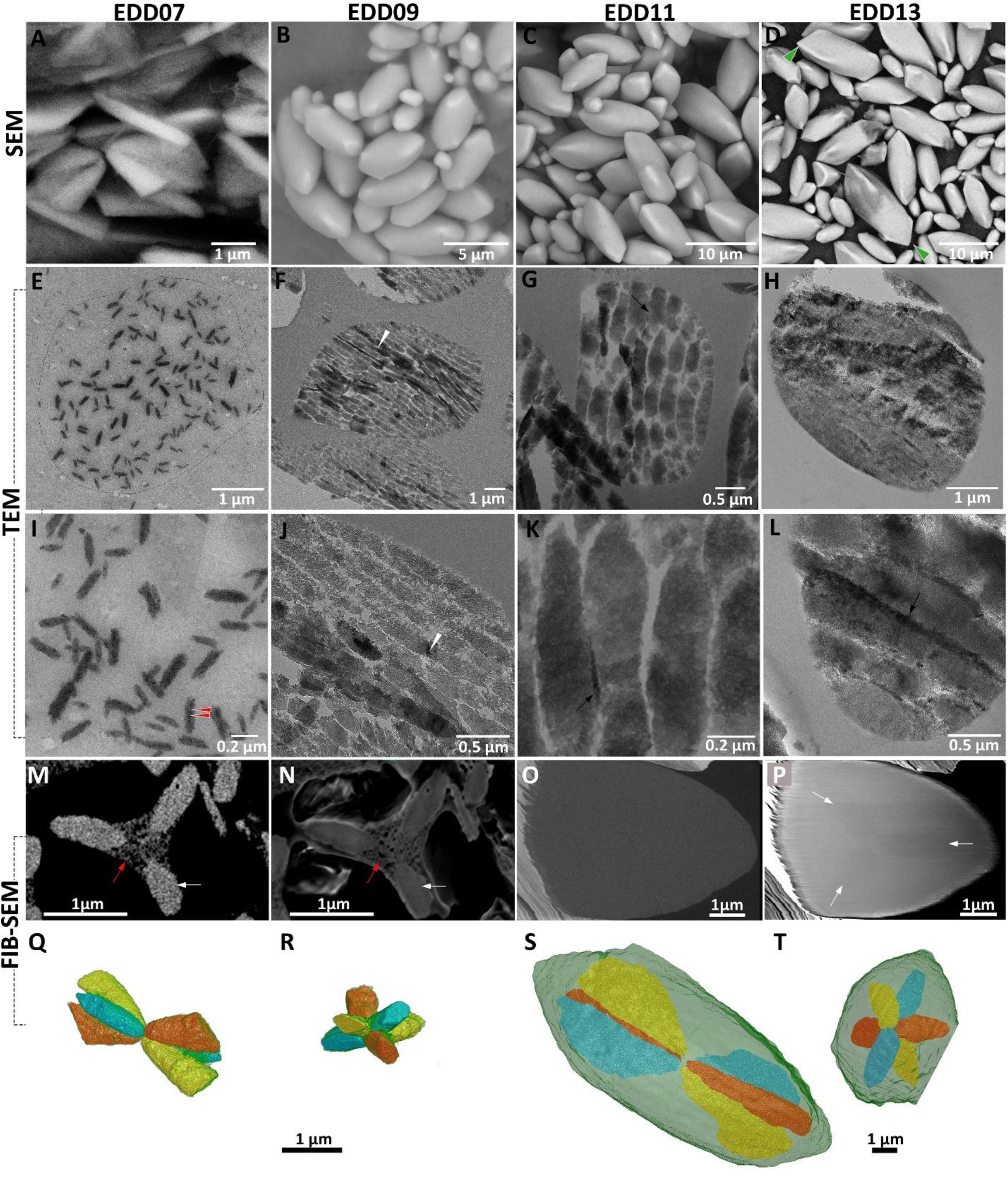
Inner structure of otoconia. (A-D) Morphology of otoconia as imaged with Scanning Electron Microscopy (SEM). (E-L) Transmitted Electron Microscopy (TEM) images: cross sections of otoconia (slices thickness 70-120 nm). EDD07 (E, I): particles with crystal domains (red arrows) randomly oriented within a template (thin black line in E indicates the boundaries of single templates). EDD09 (F, J): the particles grow and fused longitudinally (white arrowheads). EDD11 (G, K): particles grow closer and fuse transversely (black arrows). EDD13 (H, L): transverse fusion continues (black arrow) until a single particle fill the entire template (K). FIB-SEM 2D images of a single otoconium at EDD07 (M; BSE, and N; mixed InLens/SE) and at EDD13 (O; BSE, and P; mixed InLens/SE) revealed the inner organic structure of the. 3D Segmentation of the core at EDD07 (Q; longitudinal view, R; c-axis view) and at EDD13 (S; longitudinal view, T; c-axis view). Similar structure was viewed at EDD09-11 (supplementary Fig. S2.3).

TEM bright field images reveal the ultrastructural sequence of this morphological maturation (Fig.4E-L). At EDD07, dispersed and elongated mineral particles are observed within a low electron density compartment that already defines the characteristic morphology of the future otoconium (Fig. 4E, I). These particles measure ∼50 nm in width and ∼180 nm in length and frequently display multiple crystallographic subdomains (Fig. 4I; red arrowheads). At EDD09, the particles increase in size, reaching ∼150 in width and ∼450 nm in length, become preferentially aligned along their longitudinal axis (Fig. 4F), and begin to fuse with neighboring particles (Fig. 4F, J, white arrowheads). At EDD11, particle packing becomes denser and fusion extends laterally, progressively reducing the boundaries between adjacent particles (Fig. 4G, K; black arrows). At EDD13, individual particles are no longer distinguishable and the mineralized phase forms a continuous structure that occupies the entire template volume (Fig. 4H, L).

3D FIB-SEM reconstructions reveal that mature otoconia consist of a highly mineralized outer region surrounding a distinct central core that is preserved throughout development (Fig. 4M-T). Quantitative analysis demonstrates that this core occupies a remarkably constant proportion of the total otoconium volume during growth, representing approximately 15% between the EDD09 and 13. A partially mineralized otoconium identified at EDD07 provides additional insight into the sequence of mineral deposition (Fig. 4M-N). Mineralization is initially confined within the three rhombohedral facets before extending toward the central body and peripheral regions of the otoconium. This pattern is consistent with the progressive maturation observed by SEM and TEM and suggest that mineral deposition proceeds in a spatially ordered manner throughout otoconium development.

### 3.5 Structure and composition of the lagena

Selected area electron diffraction (SAED) conducted during TEM analysis demonstrates that lagena otoconia are composed of calcite. At EDD07, diffraction patterns show partially aligned polycrystalline domains rather than the continuous rings expected from randomly oriented crystallites, indicating an early degree of crystallographic organization (Fig. S2.4). From EDD09 onwards, the characteristic calcite (104) reflection is consistently observed. At EDD13, diffraction patterns acquired from different regions of a single otoconium show identical crystallographic orientations without apparent boundaries, demonstrating crystallographic continuity throughout the mineralized structure. Dark field imaging further reveals that, although most crystallites share the same orientation, some otoconia retain internal boundaries associated with slight crystallographic misorientation (Fig. S2.4C).

Synchrotron X-ray scattering analyses reveal a progressive change in the nanoscale structure of the otoconia during development (Table 2, Fig. 5A). The SAXS-derived T-like parameter, interpreted here as an apparent pore size, decreases from 5.05 ± 0.35 nm at EDD07 to 2.74 ± 0.04 nm at EDD13, accompanied by a narrower distribution. Although this parameter does not directly measure physical pores, its decrease is consistent with the reduction of the low-electron density regions between adjacent mineral particles observed by TEM as particle packing and fusion proceed. In contrast, the a- and c-lattice parameters remain stable throughout development and fall within the expected range for calcite (Table 2, Fig. 5B-C) indicating that this structural maturation occurs without detectable changes in lattice dimensions.

**Table 2.** SAXS-derived. apparent pore size and lattice parameter analysis (mean ±standard deviation) during development day (EDD).

| Development day | Apparent pore size (nm) | a-lattice parameter (nm) | c-lattice parameter (nm) |
| --- | --- | --- | --- |
| EDD07 | 5.051 ± 0.345 | 0.497 ± 0.001 | 1.703 ± 0.001 |
| EDD09 | 3.370 ± 0.277 | 0.497 ± 0.001 | 1.702 ± 0.003 |
| EDD11 | 2.873 ± 0.045 | 0.497 ± 0.001 | 1.698 ± 0.003 |
| EDD13 | 2.741 ± 0.039 | 0.498 ± 0.001 | 1.701 ± 0.004 |

**Fig 5.**
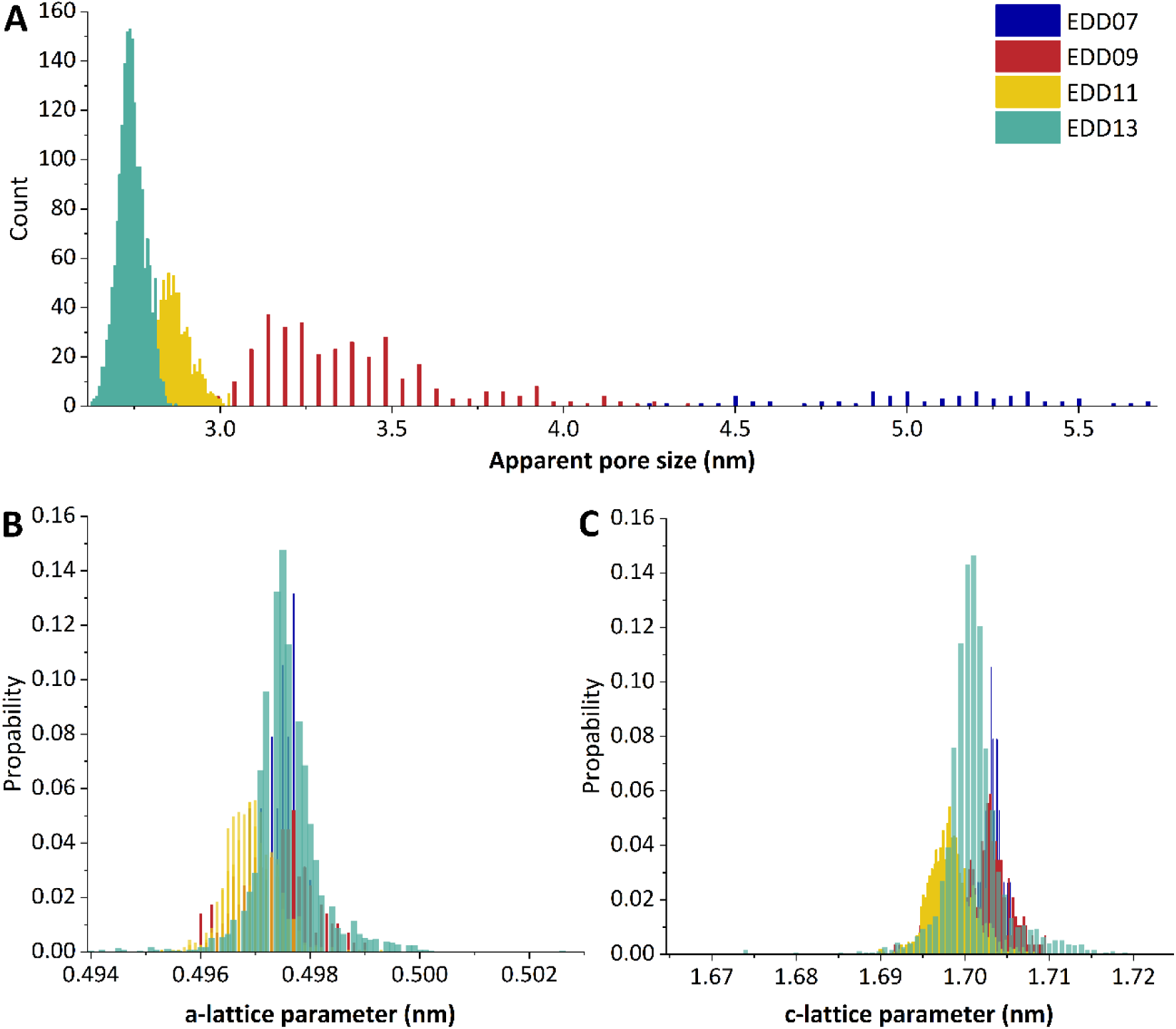
Synchrotron X-ray scattering and diffraction analyses during otoconia development. (A) Distribution of the SAXS-derived apparent pore size (represented by the T-like parameter values). (B and C) Distribution of the a-lattice parameter and the c-lattice parameters, respectively.

Raman spectroscopy and complementary elemental analyses further identify the mineral phase as magnesium substituted calcite (Fig. 6). Raman spectra acquired from three single otoconia at EDD13 show a carbonate ν_1_ peak at ∼1087 cm⁻¹ (Fig. 6A-B). Compared with geological calcite (ν_1_ ≍ 1086 cm⁻¹), this peak is shifted towards higher wavenumbers, consistent with magnesium substitution within the calcite lattice [39]. The Raman spectrum closely resembles that of quail eggshell, suggesting that both biominerals share the same magnesium calcite mineral phase. SEM EDX point measurements detect calcium, oxygen, carbon, and magnesium in all analyzed otoconia (Fig. S2.5, Table S2.2), with magnesium (Kα) representing approximately 0.6%-1.0% of the detected elements. SEM EDS lamella mapping analysis further confirms the incorporation of magnesium throughout the mineralized structure (Fig. 6C, Fig. S2.6).

**Fig 6.**
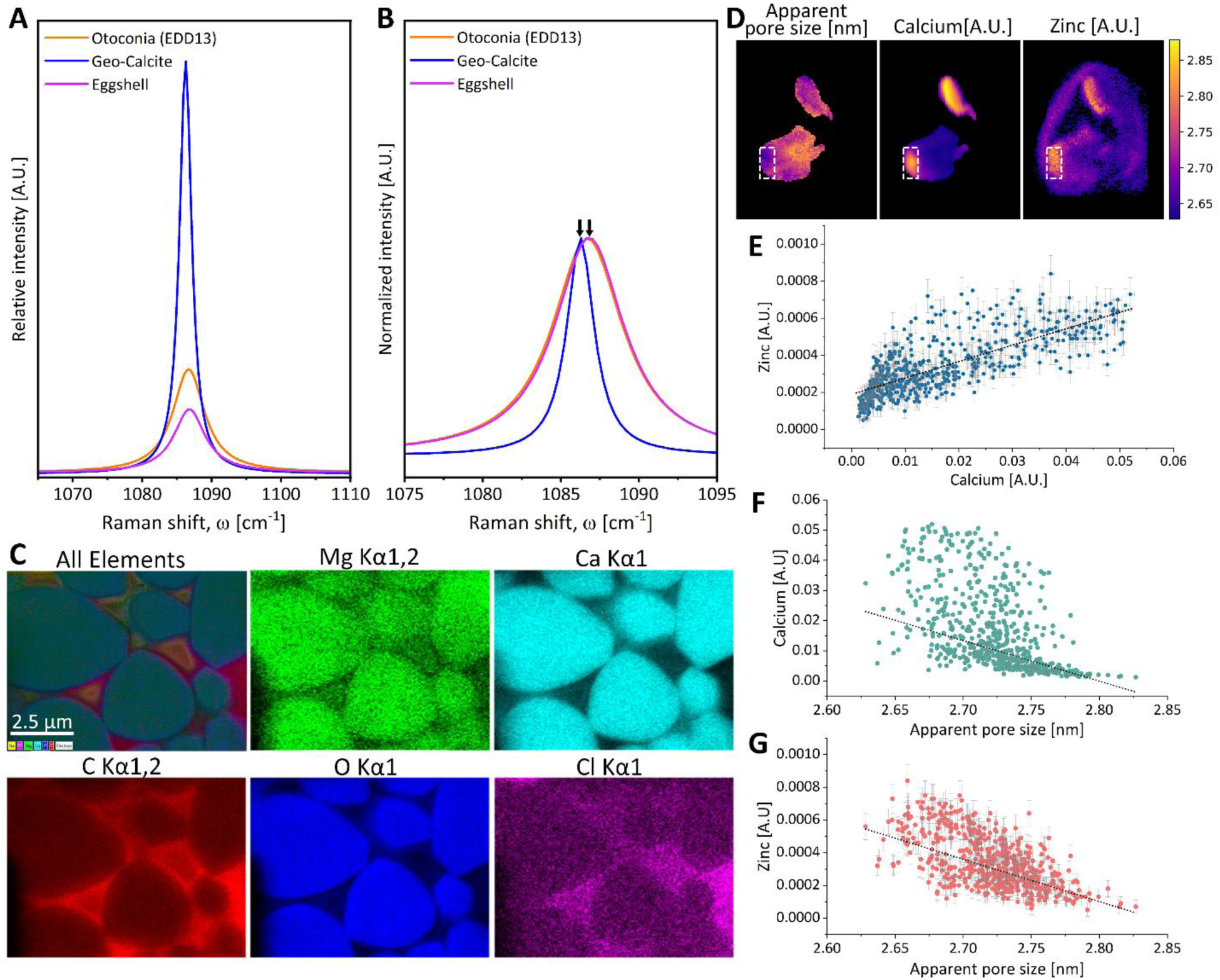
Structural and element analyses at EDD13. (A and B) Relative and normalized Raman spectra of quail otoconia, geological calcite, and quail eggshell. The carbonate ν_1_ peak of the otoconia and eggshell is shifted relative to geological calcite. (C) SEM-EDS mapping showing magnesium colocalized with calcium within the otoconia. (D) Synchrotron XRF maps of the apparent port size, calcium and zinc. Calcium is concentrated within the otoconia, whereas zinc is detected both within the mineralized structures and in the surrounding matrix. For visualization purposes, the values of calcium and zinc were rescaled to the apparent port size values (nm). Correlations (E-G) were calculated based on the values of the ROI (white rectangles in D). (E) Calcium versus zinc intensities plotted with a positive correlation slope. (F and G) Calcium and zinc verses SAXS-derived apparent pore size oplotted with a negative correlation slope. In the graphs (E-G) zinc and calcium raw recorded values are reported in arbitrary units (A.U.).

Synchrotron X-ray fluorescence (XRF) analyses detect zinc in addition to calcium within the lagena (Fig. 6D-E). Calcium and zinc intensities are positively correlated (r=0.77, *p*<0.0001) although zinc is distributed both within the otoconia and throughout the surrounding matrix. Both elemental signals are negatively correlated with the SAXS-derived apparent pore size (r(Ca)=-0.56, *p*<0.0001 and r(Zn)=-0.60, *p*<0.0001; Fig. 6F-G). These relationships indicate that regions exhibiting smaller apparent characteristic lengths generally contain higher calcium and zinc signals, but they do not establish whether zinc is incorporated into the calcite lattice or associated with the surrounding organic matrix.

## 4. Discussion

The present study provides the first multiscale characterization of otoconia development in the avian lagena and reconstructs the crystallization pathway from the earliest mineral deposits to the mature magnesium calcite crystal. By combining 3D imaging, electron microscopy, synchrotron diffraction, and spectroscopic analyses, we demonstrate that otoconia develop through progressive particle growth, alignment, and fusion within a preexisting organic compartment while maintaining a persistent central core and ultimately becoming crystallographically coherent. These observations provide new insight into the developmental assembly of calcium carbonate biomineral particles and extend previous models of otoconia formation that were largely inferred from mature specimens.

The existence of an organic compartment that regulates otoconia formation has long been proposed based on decalcification studies, growing biomimetic otoconia, and ultrastructural observations [5, 7, 11, 12, 16, 40–44]. Our observations extend this model by reconstructing the successive stages of mineralization during embryonic development. At EDD07, elongated mineral particles are already present within the organic template, defined by a low electron density compartment, that outlines the overall morphology of the future otoconium. As development proceeds, these particles progressively enlarge, align, and fuse until they become indistinguishable at EDD13. Although ultrathin sectioning may locally exaggerate the apparent spacing between mineral particles [7, 45], TEM nevertheless reveals a clear developmental reduction in the low electron density regions separating them. This ultrastructural progression is consistent with the decrease in the SAXS-derived apparent pore size. Because this parameter represents an apparent characteristic length rather than a direct measurement of physical pores, we interpret its reduction as reflecting the progressive loss of nanoscale particle/interface spacing during mineral maturation. The stable lattice parameters further indicate that this structural reorganization occurs without detectable changes in the calcite crystal lattice.

Our observations further reveal that mineral deposition within the developing otoconium follows a defined spatial sequence. Previous studies proposed that otoconia develop through sequential growth of the rhombohedral facets followed by infilling of the central body [7, 18]. Our SEM observations are consistent with this model: at EDD07, mineralization is restricted to the three rhombohedral facets, whereas the central body remains largely unmineralized. Progressive infilling of the central body is subsequently observed from EDD09 onwards, followed by continued expansion and sharpening of the rhombohedral facets during later development. These observations demonstrate that mineral deposition is spatially regulated throughout otoconium development rather than occurring uniformly across the entire structure.

3D FIB-SEM provide additional insight into this process by revealing that the mineralized shell consistently surrounds a distinct central core throughout development. Despite the substantial increase in otoconium size between EDD09 and EDD13, this compartment represents a remarkably constant proportion of the total volume, approximately 14–15%. This proportional growth suggests that the central compartment expands together with the surrounding mineral rather than being progressively replaced during maturation, supporting previous studies proposing that otoconia develop within an organic template [6, 8]. Interestingly, the core in adult guinea pig was found to be almost half of the otoconium size, which may indicate species differences in otoconium inner-structure. However, this was visually evaluated from a 2D image (deep-etching) of a freeze-fracture sample [46].

Despite their nanocomposition, mature quail otoconia behave crystallographically as single crystals. SAED analyses reveal partially aligned polycrystalline domains at EDD07, whereas from EDD09 onward the characteristic calcite diffractions are consistently detected. By EDD13, diffraction patterns acquired from different regions of individual otoconia exhibit identical crystallographic orientations without apparent grain boundaries, while dark field imaging reveals only minor internal misorientations between micro-sized domains. Together with the TEM observations, these findings indicate that early mineral particles progressively acquire crystallographic coherency during development. Similar particle mediated crystallization pathways have been proposed for several biominerals, including mouse otoconia, in which an organic matrix together with transient amorphous calcium carbonate directs the assembly of nanogranular calcite that ultimately behaves as a single crystal [44]. Likewise, mesocrystal formation has been described in numerous biomineral systems as a route toward producing highly ordered crystals through oriented particle attachment [42, 47, 48]. Our observations are consistent with such particle mediated crystallization pathway. However, because no amorphous precursor could be directly identified in the present study, the available evidence does not allow us to distinguish conclusively between classical and non-classical crystallization. Resolving this question will require future investigations combining structural characterization with thermodynamic and kinetic analyses [49].

Complementary spectroscopic and elemental analyses further show that mature quail otoconia consist of magnesium calcite. Magnesium has previously been reported in human and mouse otoconia [44, 50], where substitution of calcium by magnesium has been proposed to influence crystal morphology and mechanical properties [48, 51]. In our study, Raman spectroscopy identifies the characteristic spectral shift associated with magnesium substitution, while SEM EDS consistently detect magnesium within the mineralized otoconia. Although magnesium has been implicated in regulating crystal growth and stiffness in other biominerals, its precise contribution to otoconia formation remains unclear and will require future studies.

In addition to magnesium, synchrotron X-ray fluorescence microscopy reveals zinc within both the otoconia and the surrounding matrix. Zinc has previously been detected in vertebrate otoconia [25, 28, 52, 53], although its precise localization and role remain poorly understood. In the present study, zinc intensity is positively correlated with calcium and negatively correlated with the SAXS-derived apparent pore size. However, its broader distribution outside the mineralized structures indicates that this relationship should not be interpreted as direct evidence for zinc incorporation into the calcite lattice. Zinc may instead be associated with matrix components or zinc-dependent enzymes involved in calcium carbonate formation, including carbonic anhydrase [4, 52]. Zinc is recognized as an important regulator of biomineralization in numerous mineralized tissues, where it can influence crystal nucleation, crystal growth, and the activity of matrix associated proteins [54]. However, determining whether zinc contributes directly to crystal formation or reflects enzymatic activity in the surrounding matrix will require higher resolution elemental and molecular localization.

Beyond the ultrastructural observations, our quantitative analyses provide new insight into the developmental growth and spatial organization of the avian lagena. Mineralization is already established at EDD07, before detectable mineralization of the surrounding cranial bones, indicating that vestibular biomineralization is initiated very early during embryogenesis. This agrees with previous developmental studies showing that formation of the vestibular sensory organs precedes skeletal mineralization [18, 55, 56]. Although the exact onset of mineral deposition could not be determined here, the presence of numerous mineralized otoconia at EDD07 suggests that crystallization is initiated shortly beforehand.

The most pronounced increase in otoconia number and mineralized volume occurs between EDD07 and EDD09, consistent with the rapid developmental expansion previously described in the avian saccule [18] and during early human vestibular development [57]. However, our 3D quantitative analyses demonstrate that total mineralized volume increases proportionally more than otoconia number, indicating that enlargement of existing otoconia contributes more to overall mineral accumulation than the continuous formation of new crystals. At the same time, small otoconia remain present throughout development, suggesting that nucleation and crystal growth occur simultaneously within the otolithic membrane. To our knowledge, this represents the first quantitative evidence of these growth dynamics in the avian lagena.

3D reconstructions reveal a highly organized spatial arrangement of otoconia. Larger crystals are preferentially located near the periphery of the lagena, whereas smaller otoconia become concentrated above the striola during development. Comparable regional variations have been reported in the utricle and saccule of birds and mammals, including the “snowdrift” region described in guinea pig vestibular organs [5, 10, 18, 58, 59]. Although a distinct snowdrift morphology was not observed in the quail lagena, the accumulation of smaller otoconia above the striola suggests that spatial regulation of crystal size is a conserved characteristic of vertebrate vestibular organs. Previous study proposed that this regional specialization may reflect differences in mechanical loading or sensory innervation across the macula [18]. While the mechanisms responsible for establishing this organization remain unknown, our observations indicate that otoconia growth is tightly coordinated with maturation of the sensory epithelium. The overall morphology of mature quail otoconia is comparable to that described in the utricle and saccule of birds and mammals, including the occurrence of occasional large (“giant”) otoconia [10, 15, 44, 51]. Rather than revealing a unique crystal morphology, the avian lagena appears to share the same fundamental biomineralization strategy as the other vestibular organs. The distinctive characteristics of the lagena therefore likely arise from the 3D organization of the organ itself rather than from differences in the ultrastructure or composition of individual otoconia. This observation has important implications for understanding the physiological role of the avian lagena. Although auditory and magnetoreceptive functions have been proposed [25–28], electrophysiological studies have shown that lagena afferents do not participate in auditory processing [21–23], and recent evidence suggests that magnetic sensing in pigeons is primarily mediated by the semicircular canals [60]. Our findings instead support a vestibular function. The internal architecture of quail otoconia closely resembles that of artificial otoconia shown by particle dynamics simulations to optimize responses to gravity and linear acceleration through their asymmetric mass distribution [61]. Together with their magnesium calcite composition, these features suggest that lagena otoconia are well suited for gravity sensing rather than representing a specialized biomineral.

If the avian lagena fulfills a distinct functional role, our results suggest that this specialization is more likely to arise from the geometry of the organ than from the properties of individual otoconia. Unlike the utricle and saccule, which are organized largely within a single plane, the lagena extends through 3D and contains multiple curvatures. Such geometry has been proposed to broaden the range of gravitational and inertial stimuli detected by the sensory epithelium [62, 63]. Because birds rely on rapid and continuous head movements to stabilize gaze during locomotion and flight through the vestibulo ocular reflex [64, 65], we speculate that the lagena may provide complementary information on gravity and linear acceleration during these movements. Rather than acting as a dedicated navigation or magnetoreceptive organ, the avian lagena may therefore enhance vestibular performance through its unique 3D morphology.

## 5. Conclusion

This study provides the first multiscale characterization of otoconia development in the avian lagena. We demonstrate that lagena otoconia develop through progressive particle growth, alignment, and fusion within a persistent organic compartment, ultimately forming crystallographically coherent magnesium calcite crystals. Our results further reveal that otoconia growth is spatially regulated within the developing lagena and follows the same fundamental biomineralization strategy as other vertebrate vestibular organs. Our results establish a developmental model of avian otoconia formation and provide a structural foundation for future studies investigating the relationship between vestibular biomineralization and function.

## CRediT authorship contribution statement

**Einat Kedar:** Conceptualization, Methodology, Investigation, Formal analysis, Validation, Visualization, Writing – original draft, Writing – review & editing. **Jia Hui Lim:** Methodology, Formal analysis, Writing – review & editing. **Ernesto Scoppola:** Methodology, Formal analysis, Writing – review & editing. **Peter Fratzl:** Conceptualization, Data curation, Writing – review & editing. **Shahrouz Amini:** Methodology, Formal analysis, Investigation, Writing – review & editing. **Emeline Raguin:** Conceptualization, Methodology, Formal analysis, Supervision, Writing – review & editing.

## Declaration of competing interest

The authors declare that they have no known competing financial interests or personal relationships that could have appeared to influence the work reported in this paper.

## Acknowledgments

We thank Susann Weichold for technical support with the ultrathin sections and SEM imaging, Jeannette Steffen for histology staining and Heike Runge for technical support with the FIB-SEM (all in the Department of Biomaterials at the Max Planck Institute of Colloids and Interfaces). We thank Yu Ogawa for advising with the TEM imaging and diffraction analysis (Department of Sustainable and Bio-inspired Materials at the Max Planck Institute of Colloids and Interfaces).

## Notes

### Competing Interest Statement

The authors have declared no competing interest.

