## Supplementary for "Crystallization of magnesium calcite otoconia in the inner ear of the developing quail"

**Supplementary S1**

### S1.1 Otoconia extraction

Under a dissection microscope (Vision Engineering MC-Uni Mantis Compact Universal Stereo Microscope), the lagena otoconia was extracted after cutting the head in half along the dorsal midline of the skull. After cutting the skull to half at midline, the brain was removed to expose the inner ear (Fig. S1.1). Then the lagena (L) was recognize at the end of the basilar papilla and gently removed. U: utricle, S: saccule.

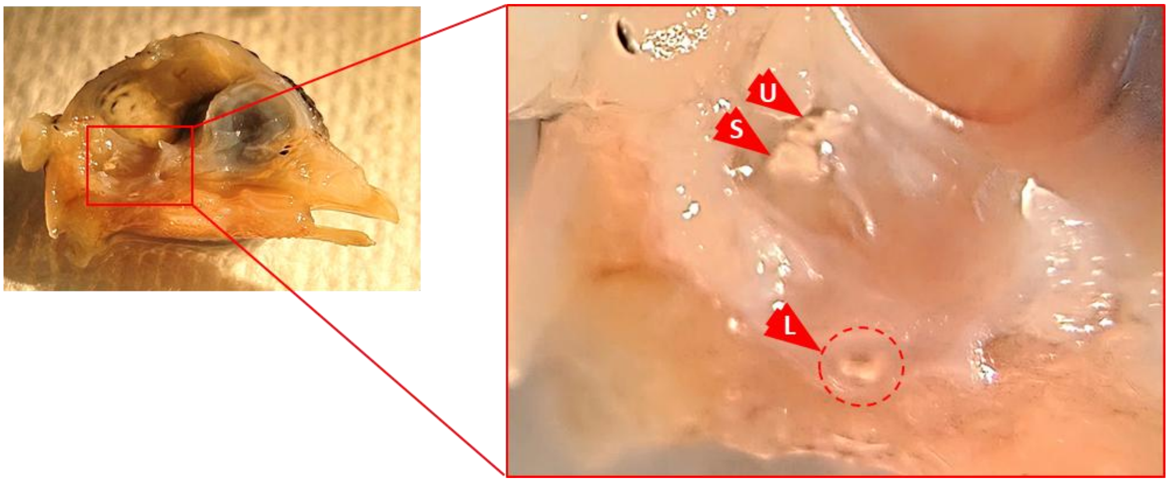


**Fig. S1.1 Dissection microscope view:** Left inner ear and otoliths: U: utricle, S: saccule, L: lagena.

#### S1.2 Lamella preparation

Lamellae were prepared using a ZEISS Crossbeam 550 FIB-SEM (Carl Zeiss Microscopy GmbH, Germany) equipped with a gallium ion source operated at 30 kV and an OmniProbe 350 cryo-micromanipulator (Oxford Instruments, United Kingdom). Prior to trench milling, a protective platinum (Pt) layer approximately 1.5 µm thick was deposited over the region of interest. Coarse and fine trench milling was performed at ion beam currents ranging from 3 to 15 nA, producing a lamella with approximate dimensions of 20 µm (length) × 12 µm (height) × 2 µm (width). The lamella was subsequently lifted using the OmniProbe micromanipulator needle and transferred to a lift-out half-moon grid. Final polishing was carried out from the front face using a 50 pA ion beam, yielding a final section thickness of approximately 1.5 µm.

**Supplementary S2**

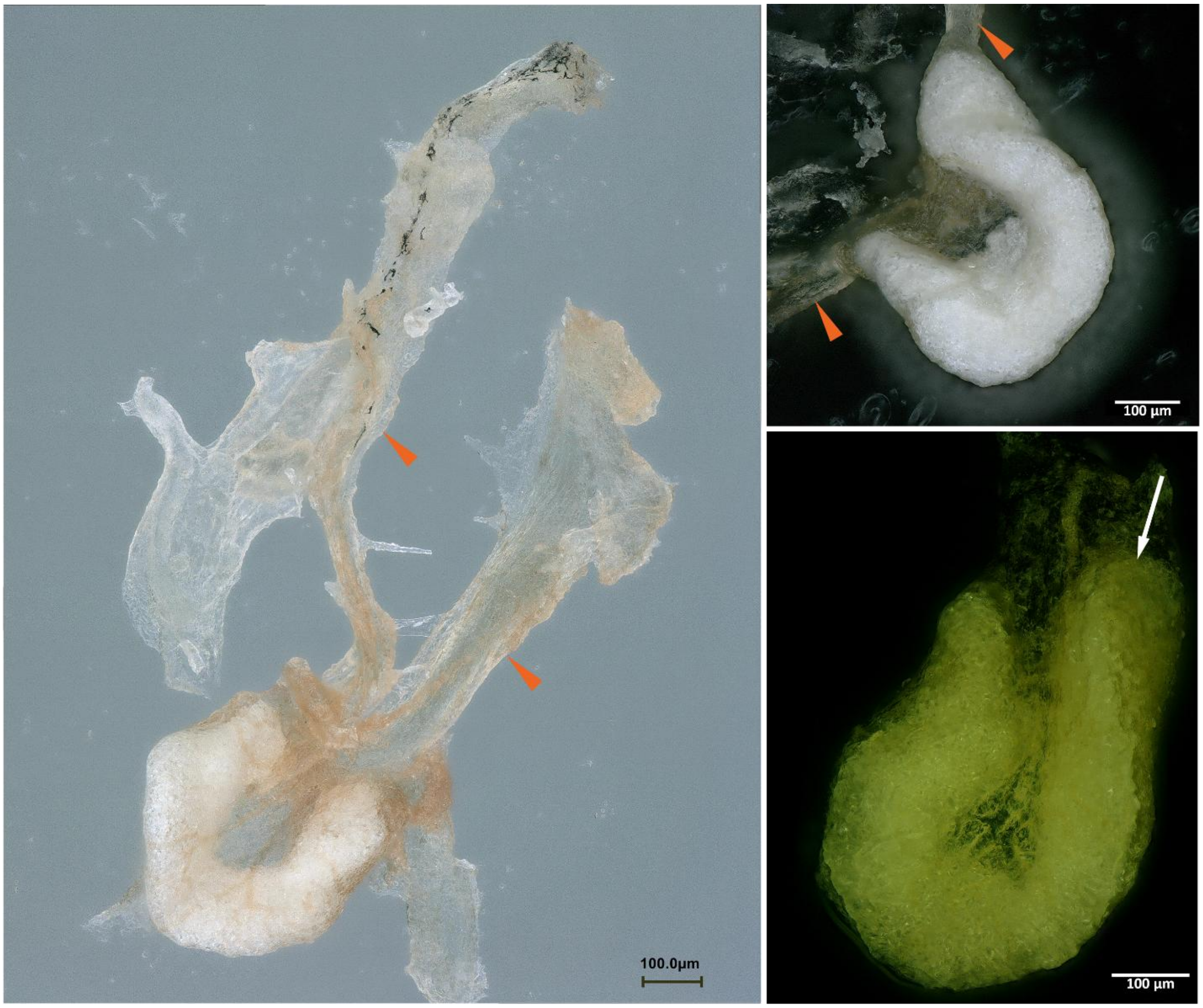


**Fig. S2.1**. Light microscopy images of EDD10 on glass slide (A), EDD11 on a black carbon tape (B) and EDD13 on a black carbon tape (C). During development, the otoconia grow within the boundaries of the otolith membrane. Notice the membrane extensions to each side of the lagena (orange arrows). In mature otoconia, one arm is longer than the other (C, white arrow).

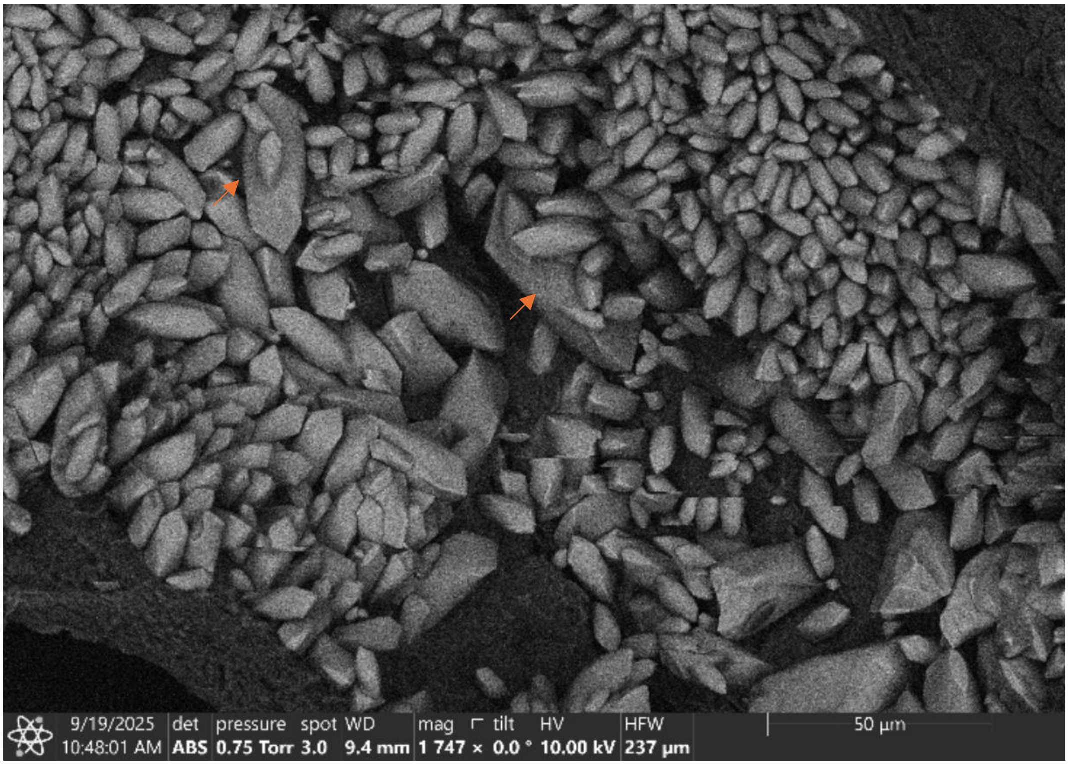


**Fig. S2.2. Giant otoconia.** SEM image of EDD13. Arrows point to otoconia that are >30µm.

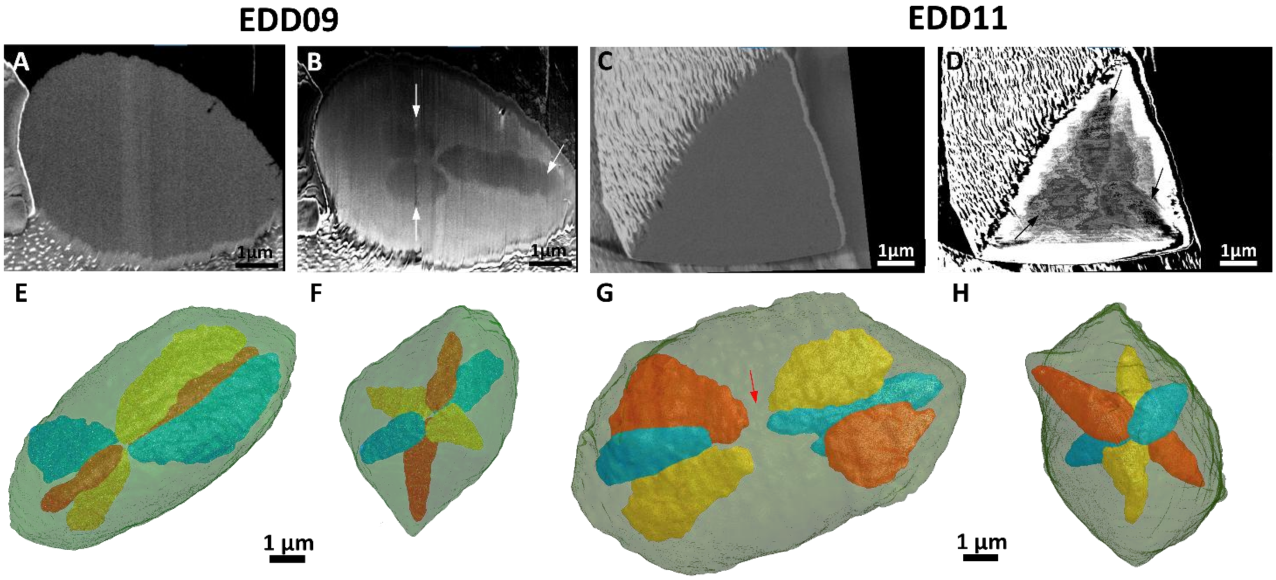


**Fig. S2.3** FIB-SEM images at EDD09 and EDD11 show similar architecture like EDD13 (see manuscript Fig. 4). 2D BSE (A) and mixed InLens/SE (B) at EDD09, and BSE (C) and mixed InLens/SE (D) at EDD11. White and black arrow show the organic core in appear darker in the mixed images. The core architecture is presented in 3D at EDD09 (E-F) and EDD11 (G-F). Milled otoconium at EDD11 was disturbed due to artifact in the image, therefore a gap was form in the core construction and segmentation (red arrow).

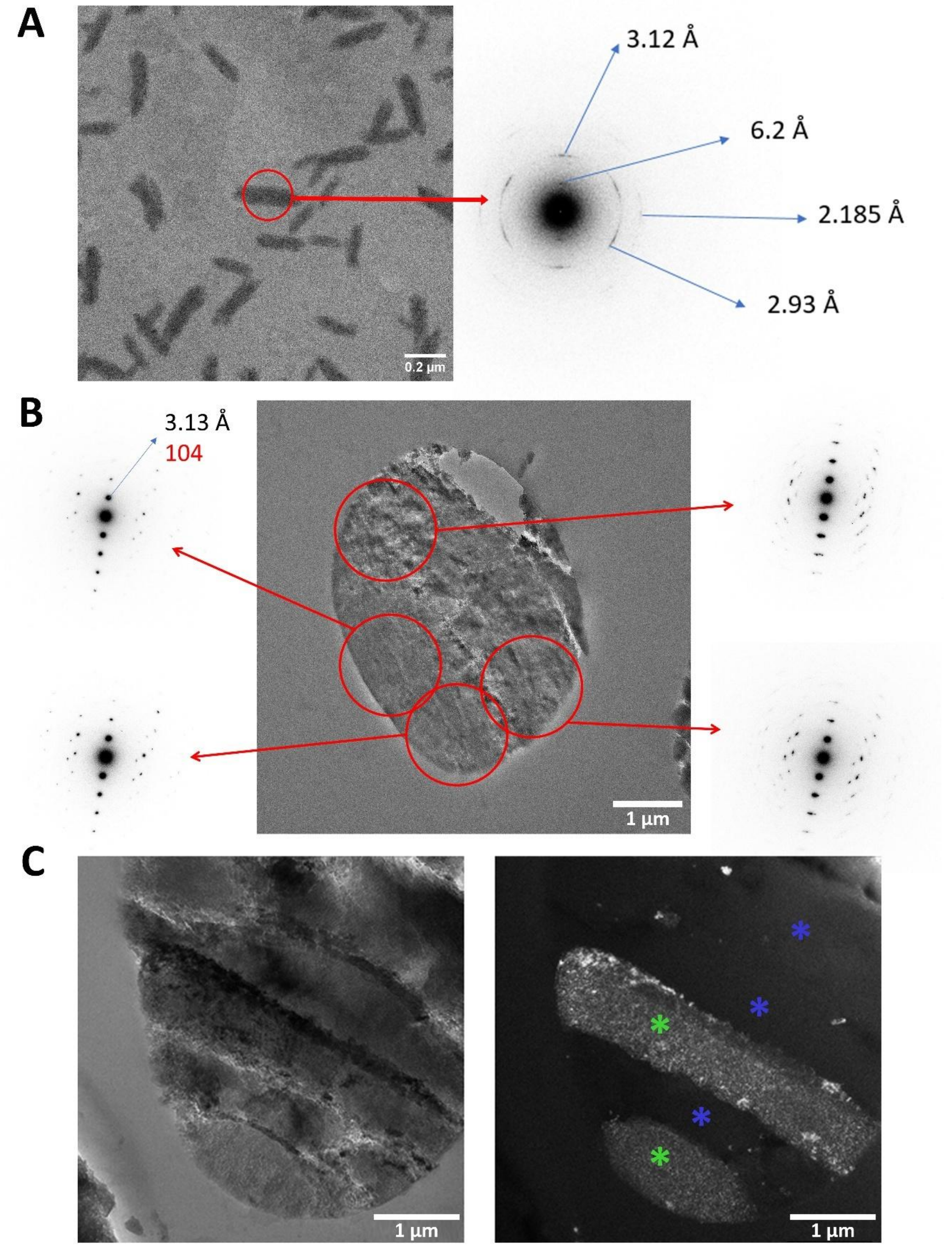


**Fig. S2.4. Selected area electron diffraction (SAED) pattern** of A. EDD07 120nm thickness and B. EDD13. C. right- bright field image (normal image), left- dark field image (normal image) blue Asterix: crystals that do not share the same orientation, green Asterix: crystals that share the exact orientation.

**Table S2.1.** Apparent pore size and lattice analysis, by development day (EDD).

| | N | Mean ±standard deviation | SE of mean | Lower 95% CI of Mean | Upper 95% CI of Mean | Skewness | Range | Median |
|---|---|---|---|---|---|---|---|---|
| **Apparent pore size** | | | | | | | | |
| EDD07 | 73 | 5.051 ±0.345 | 0.040 | 4.972 | 5.132 | -0.347 | 4.25-5.702 | 5.102 |
| EDD09 | 347 | 3.370 ±0.277 | 0.015 | 3.340 | 3.399 | 1.017 | 2.7-4.362 | 3.335 |
| EDD11 | 702 | 2.873 ±0.045 | 0.002 | 2.870 | 2.877 | 0.810 | 2.789-3.025 | 2.865 |
| EDD13 | 1817 | 2.741 ±0.039 | 9.12E-04 | 2.740 | 2.743 | 0.070 | 2.626-2.871 | 2.737 |
| **a-Lattice** | | | | | | | | |
| EDD07 | 38 | 0.497 ±0.001 | 5.02465E-5 | 0.497 | 0.497 | -0.919 | 0.497-0.498 | 0.497 |
| EDD09 | 289 | 0.497 ±0.001 | 3.64797E-5 | 0.497 | 0.497 | -0.126 | 0.496-0.499 | 0.497 |
| EDD11 | 1147 | 0.497 ±0.001 | 1.33151E-5 | 0.497 | 0.497 | 0.2965 | 0.495-0.499 | 0.497 |
| EDD13 | 2728 | 0.498 ±0.001 | 1.18E-5 | 0.498 | 0.498 | 0.309 | 0.494-0.503 | 0.497 |
| **c- Lattice** | | | | | | | | |
| EDD07 | 38 | 1.703 ±0.001 | 0.001 | 0.000 | 1.703 | 1.704 | 0.249-1.703 | 1.702 |
| EDD09 | 289 | 1.702 ±0.003 | 0.003 | 0.000 | 1.702 | 1.703 | -0.527-1.703 | 1.692 |
| EDD11 | 1147 | 1.698 ±0.003 | 0.003 | 0.000 | 1.698 | 1.699 | 0.395-1.698 | 1.690 |
| EDD13 | 2728 | 1.701 ±0.004 | 0.004 | 0.000 | 1.701 | 1.701 | -0.555-1.701 | 1.670 |

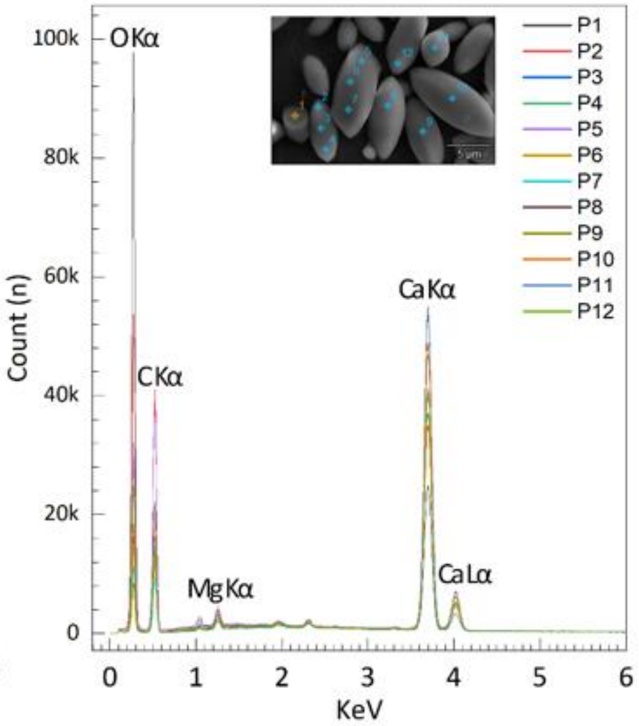


**Fig. S2.5. SEM EDX spectrum of otoconia collected as a series of point measurements at EDD13 (extracted otoconia, 10nm carbon coating).**

**Table S2.2. Elemental analysis using SEM EDX in A) Weight percentage and B) Atomic percentage.**

A. Weight %

| | ***C*** | ***O*** | ***Mg*** | ***Al*** | ***Si*** | ***P*** | ***S*** | ***K*** | ***Ca*** | ***Cu*** |
|---|---|---|---|---|---|---|---|---|---|---|
| ***pt1*** | 47.606 | 31.631 | 0.564 | 0.044 | 0.065 | 0.151 | 0.562 | 0.081 | 19.295 | |
| ***pt2*** | 27.836 | 46.437 | 0.709 | 0.074 | | 0.150 | 0.389 | 0.084 | 24.321 | |
| ***pt3*** | 16.023 | 35.906 | 1.047 | 0.043 | 0.078 | | 0.552 | 0.185 | 45.754 | 0.412 |
| ***pt4*** | 14.231 | 33.666 | 0.986 | | 0.065 | | 0.538 | 0.177 | 50.337 | |
| ***pt5*** | 17.369 | 47.592 | 0.863 | 0.067 | | | 0.272 | 0.129 | 33.707 | |
| ***pt6*** | 13.806 | 33.305 | 0.893 | 0.059 | | | 0.504 | 0.181 | 51.252 | |
| ***pt7*** | 12.166 | 32.032 | 0.855 | | 0.084 | | 0.580 | 0.160 | 54.124 | |
| ***pt8*** | 12.882 | 32.285 | 0.910 | 0.040 | | | 0.347 | 0.146 | 53.391 | |
| ***pt9*** | 7.793 | 35.106 | 0.835 | | | | 0.386 | 0.154 | 55.727 | |
| ***pt10*** | 13.995 | 38.504 | 0.986 | 0.046 | | | 0.400 | 0.167 | 45.902 | |
| ***pt11*** | 18.575 | 32.626 | 0.984 | 0.039 | 0.063 | | 0.252 | 0.139 | 47.321 | |
| ***pt12*** | 20.787 | 35.675 | 0.566 | 0.051 | 0.084 | | 0.559 | 0.147 | 42.131 | |

B. Atom %

| | ***C*** | ***O*** | ***Mg*** | ***Al*** | ***Si*** | ***P*** | ***S*** | ***K*** | ***Ca*** | ***Cu*** |
|---|---|---|---|---|---|---|---|---|---|---|
| ***pt1*** | 61.226 | 30.539 | 0.359 | 0.025 | 0.036 | 0.075 | 0.271 | 0.032 | 7.437 | |
| ***pt2*** | 39.429 | 49.379 | 0.496 | 0.047 | | 0.083 | 0.206 | 0.036 | 10.324 | |
| ***pt3*** | 27.817 | 46.796 | 0.898 | 0.033 | 0.058 | | 0.359 | 0.099 | 23.804 | 0.135 |
| ***pt4*** | 25.706 | 45.653 | 0.880 | | 0.051 | | 0.364 | 0.098 | 27.248 | |
| ***pt5*** | 27.226 | 56.003 | 0.669 | 0.047 | | | 0.160 | 0.062 | 15.833 | |
| ***pt6*** | 25.157 | 45.559 | 0.805 | 0.048 | | | 0.344 | 0.102 | 27.987 | |
| ***pt7*** | 22.887 | 45.237 | 0.795 | | 0.067 | | 0.409 | 0.093 | 30.512 | |
| ***pt8*** | 23.961 | 45.082 | 0.837 | 0.033 | | | 0.242 | 0.083 | 29.761 | |
| ***pt9*** | 15.146 | 51.222 | 0.802 | | | | 0.281 | 0.092 | 32.457 | |
| ***pt10*** | 24.396 | 50.389 | 0.849 | 0.036 | | | 0.261 | 0.089 | 23.979 | |
| ***pt11*** | 32.072 | 42.290 | 0.840 | 0.030 | 0.046 | | 0.163 | 0.074 | 24.485 | |
| ***pt12*** | 34.197 | 44.058 | 0.460 | 0.037 | 0.059 | | 0.345 | 0.074 | 20.770 | |

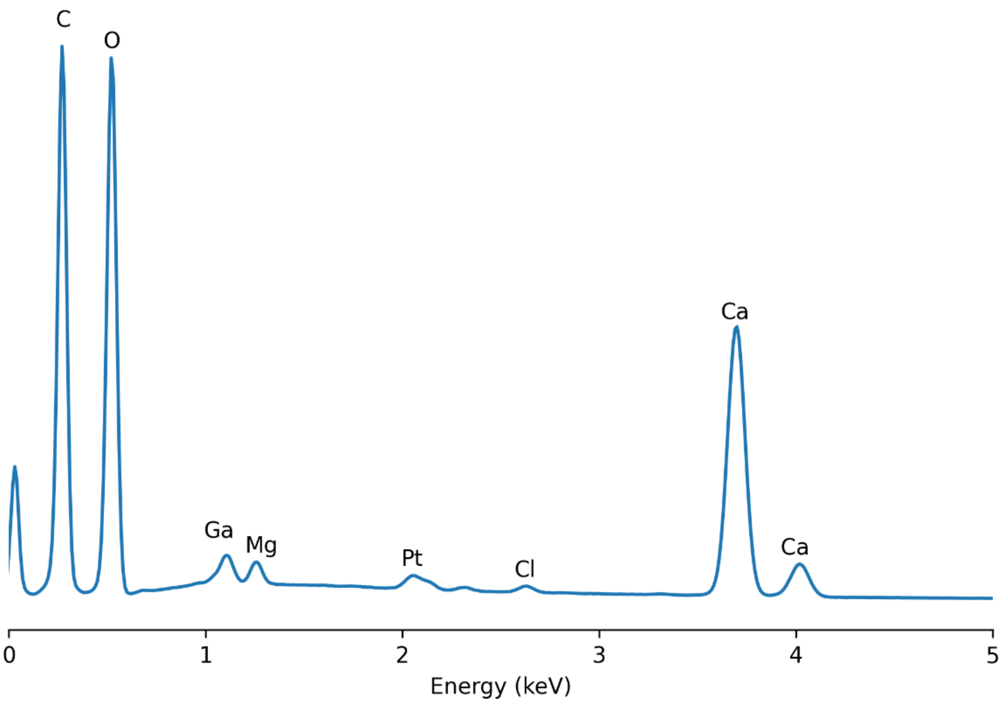


**Fig. S2.6. SEM EDS spectrum of otoconia at EDD13** (resin embedded lamella). Mapping analysis revealed the presence of carbon, oxidant, gallium (from the gallium source), magnesium, platinum, chlorine, and calcium elements.
